# ShapeProt: Top-down Protein Design with 3D Protein Shape Generative Model

**DOI:** 10.1101/2023.12.03.567710

**Authors:** Youhan Lee, Jaehoon Kim

## Abstract

With the fact that protein functionality is tied to its structure and shape, a protein design paradigm of generating proteins tailored to specific shape contexts has been utilized for various biological applications. Recently, researchers have shown that top-down strategies are possible with the aid of deep learning for the shape-conditioned design. However, state-of-the-art models have limitations because they do not fully consider the geometric and chemical constraints of the entire shape. In response, we propose ShapeProt, a pioneering end-to-end protein design framework that directly generates protein surfaces and generate sequences with considering the entire nature of the generated shapes. ShapeProt distinguishes itself from current protein deep learning models that primarily handle sequence or structure data because ShapeProt directly handles surfaces. ShapeProt framework employs mask-based inpainting and conditioning to generate diverse shapes at the desired location, and these shapes are then translated into sequences using a shape-conditioned language model. Drawing upon various experimental results, we first prove the feasibility of generative design directly on the three-dimensional molecular surfaces beyond sequences and structures.

## 1 Introduction

Proteins have a vital function in coordinating many biological processes through their interactions with other biomolecules. Hence, protein design, the procedure of crafting protein sequences for specific objectives, holds significant significance in diverse domains [1, 2, 3, 4, 5, 6, 7]. Since the functionality of proteins is intricately defined by the overall structure and the molecular surface of protein function sites, protein sequence design can be regarded as the design of protein shapes. Building upon these principles, numerous researchers have endeavored to design proteins that fit specific shapes, a critical requirement for applications such as vaccine creation. Fragment-based bottom-up methodologies have generally dominated the discipline, but recent improvements have brought a new paradigm called the top-down approach with the aid of deep learning. For instance, Lutc et al.[8] utilized reinforcement learning methods to assemble protein based on global geometric constraints represented as scalar values. On the other hand, Chroma [9] suggested a sampling method for generating proteins that closely resemble the morphologies of the provided point clouds. Although these methods have made significant advances by introducing innovative design paradigms, they still need to be revised to comprehensively consider all aspects of protein shape, namely, chemical and geometric complementarity determined by participating amino acids [10, 11].

In recent years, we have witnessed the remarkable success of artificial intelligence, and notably, data-driven generative AI is transforming and influencing various fields and our lives through chatbots [12, 13], image generation [14, 15, 16], and video synthesis [17, 18]. The field of protein design has also benefited significantly from the generative AI in recent years [19]. As essential data modalities in the realm of proteins are sequences and structures, many studies have introduced corresponding generative AI techniques and shown promising results. Specifically, protein sequence databases comprise hundreds of millions of sequences, encompassing various species and diverse families. By applying well-established pre-training, such as masked language modeling (MLM) [20], causal language modeling (CLM) [21], and Fill-in-Middle language modeling (FIM) [22] on the sequence database, several protein language models have been developed [23, 24, 25, 26]. These models have been utilized in diverse protein sequence design tasks, leading to substantial advancements with improved efficacy compared to previous approaches. For instance, the ESM [23] model is primarily used for high-quality representation learning and is further expanded for sequence design in combination with conditional masked language modeling [27]. Moreover, models like ProGen [25] and ProtGPT2 [24], which employed causal language modeling, utilized to produce sequences in response to a specific biological context. In addition to this, SaProt [28] suggested a variant of MLM on mixed sequences obtained by mixing amino acids sequence and structure sequence calculated by FoldSeek [29]. On the other hand, many researchers have been interested in structural context-based sequence generation using characterized and predicted structure data. In particular, the deep-learning-based inverse folding, approach for tackling fixed backbone design has demonstrated encouraging outcomes [30, 31, 32]. In the method, a structure encoder is used to obtain the structural context from the given backbone, and this context is translated into a sequence through captioning or sampling. This approach appends suitable amino acids continuously conditioned on the fixed backbone during generation, ensuring the preservation of a stable structure while allowing for the sampling of diverse sequences. These approaches were empirically proven more effective than traditional energy function-based design methods [19]. Furthermore, techniques that simultaneously model and generate structure and sequence have been proposed by primarily employing iterative refinement [33]. However, interestingly, the generative modeling directly on three-dimensional protein surfaces have been unexplored. We imagined that generating protein surfaces with various geometrical and chemical attributes will flourish the resources of templates for the top-down based protein design.

In the realm of generative AI, an important branch gaining a reputation is 3D generative modeling, which is actively explored for its applications in neural rendering [34, 35, 36]. Within 3D generative modeling, 3D assets are represented in various data formats, such as point clouds, meshes, voxels, or implicit neural representations. Generation is thereafter performed based on a model that captures the data distribution. Notably, the incorporation of conditioning techniques has demonstrated the ability for user-controllable generation [37, 38]. Prompted by these achievements, we contemplated the prospect of training a generative model to (i) grasp the intricacies of protein surfaces, (ii) proficiently generate diverse surfaces, and (iii) employ these newfound shapes as conditioning templates to guide top-down protein sequence design.

In this work, we propose ShapeProt, a novel and new type of protein generative AI model that directly learns proteins’ geometrical and chemical surfaces and can generate surfaces based on the learned distribution. ShapeProt utilizes a 3D Vector-Quantized Variational Autoencoder (3D-VQVAE) [39] and latent diffusion models (LDM) [16]. Additionally, we developed a surface-conditioned language model to generate sequences that match the generated shapes. Notably, we showed that ShapeProt is able to perform molecular surface surgery where the target local protein shape is replaced with a generated shape possessing the desired properties through inpainting with classifier-free guidance [40].

## 2 Results

### 2.1 Voxelized Protein Surface Database

In this study, we adopted the molecular surface interaction fingerprinting (MaSIF) representation for characterizing the protein surfaces [41]. Within the MaSIF, the initial step is to obtain a molecular shape mesh based on the Connolly surface [42, 43]. For each mesh vertex, three features are calculated: hydrophobicity, electronic charge, and hydrogen bond potential. Our goal was to develop a generative model capable of processing the three-dimensional shapes of proteins and their associated colors, i.e., chemical properties. To achieve this goal, we employed the Signed distance function (SDF)-based 3D deep learning modeling strategy, which have recently achieved significant achievements on representation and generation of 3D objects [44, 45].

The SDF, as illustrated in Figure 1 (a), refers to the distance from a certain query point to the closest point on the surface of the object being modeled. According to this definition, the SDFs of points on the surface are zero, whereas the rest are expressed as signed distances. In this work, we depicted the inner SDF as negative and the outer surface as positive. By utilizing this SDF form, we are able to estimate an implicit surface. Significantly, the 3D asset represented using SDF is readily converted into a 3D mesh by employing the well-defined Marching Cubes algorithm [46]. Indeed, the concept of SDF have been utilized in molecular visualization [47].

**Figure 1:**
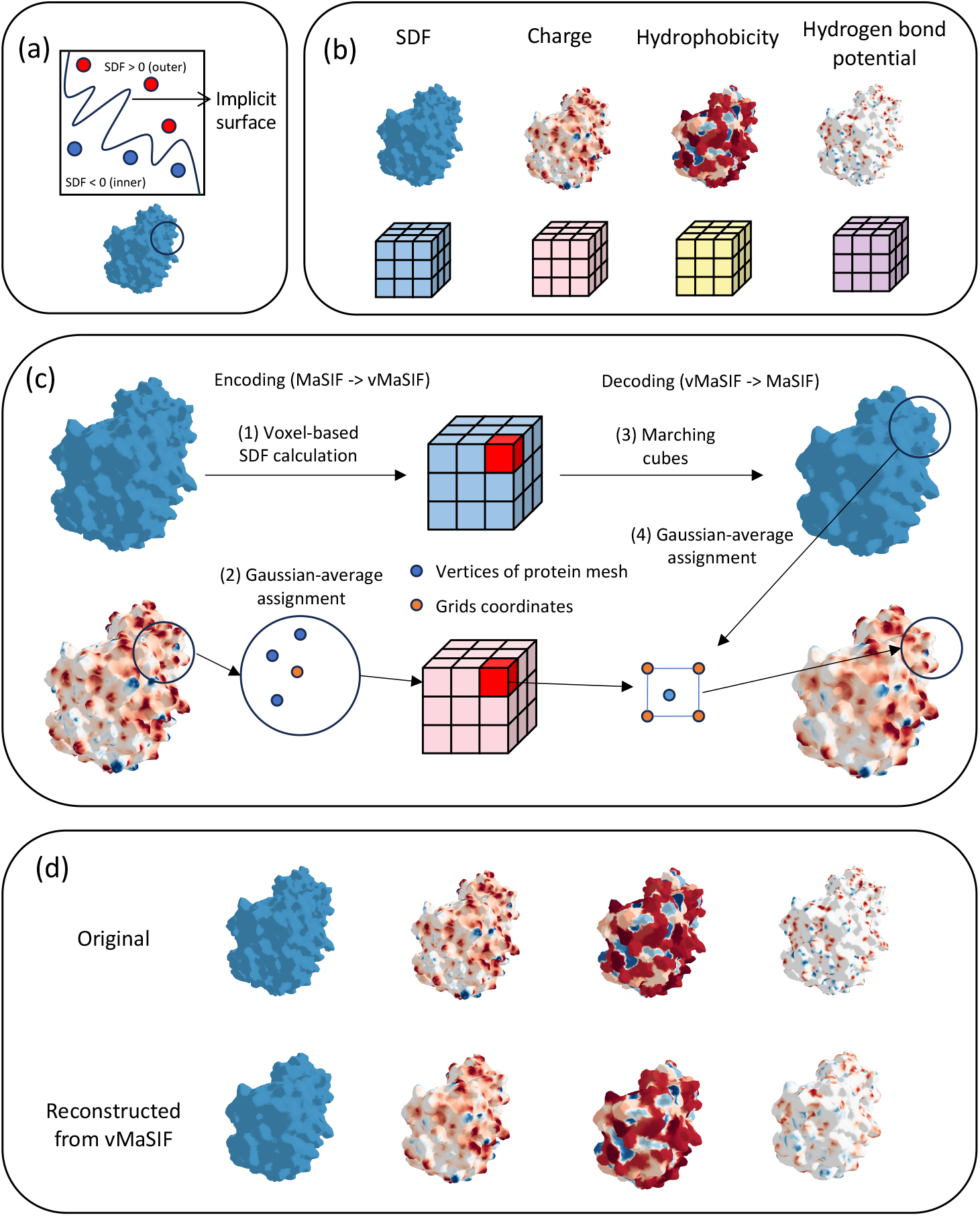
(a), a conceptual illustration of signed distance functions for protein surfaces. (b), MaSIF based four representations. We choose SDF representation for geometric properties. (c), Encoding and decoding process for generating voxelized MaSIF (vMaSIF). (d), Some examples reconstructed from our voxelized MaSIF.

An arbitrary query point set is required to calculate the Signed Distance Function (SDF). So, we utilized a 3D voxelized SDF where grid coordinates represent pre-defined groupings of points. This method is in line with the objective of our research, which is the generation and modification of 3D protein surfaces, for two reasons. Firstly, voxelized input allows a straightforward extension of multiple features. Our input includes the SDF, which represents the shape, and the surface colors describing three surface chemical characteristics (See Figure 1 (b)). By adding three channels representing chemical features to the SDF channel-wise, we effectively converted MaSIF into a four-channeled voxel input. Second, multi-channel voxel can be effectively modeled by employing 3D Convolutional Neural Networks (3D-CNN) [48]. Furthermore, the coordinate system of the latent grid voxel acquired by 3D-CNN represents the real-space coordinate system, allowing us to employ it for inpainting effectively. This topic will be elaborated extensively in the following results.

The our data preparation follows: First, we center the computed molecular mesh. Subsequently, we obtain the SDF voxel grid using distances between pre-defined grid coordinates and vertices of the surface mesh. The chemical features for coordinates of the voxel grid are calculated based on the Gaussian-weighted average of charge features from the top K nearest vertices according to distances. Converting the voxel grid back to a molecular mesh starts with calculating the molecular mesh using marching cubes. Then, we similarly take the Gaussian-weighted average of chemical features from the top K nearest voxel grid points and assign the features to the vertices. Figure 1 (c) illustrates the original and corresponding surface data after the our grid voxel generation and reconstruction. As depicted in Figure 1 (d), the details are well-preserved. We prepared about 440,000 grid voxels from the PDB database. An appropriate size of a voxel grid must be defined to depict 3D shapes, as large grids incur high computational costs. We determined a voxel size 96 through preliminary tests, which is computationally affordable while maintaining high fidelity (Supplementary). We abbreviate voxelized MaSIF as vMaSIF in the following.

### 2.2 3D Vector-Quantized Variational Autoencoder for Protein Surface

**Table 1:**
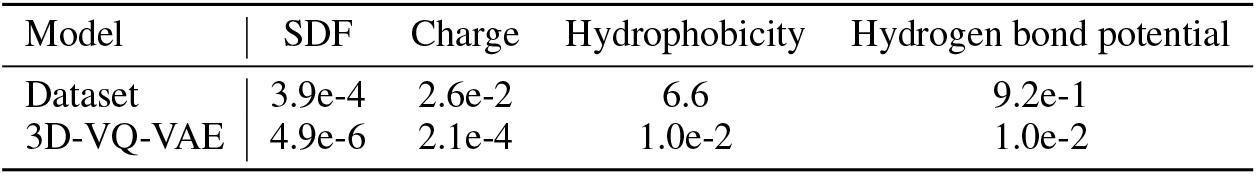
Mean squared errors between vMaSIF. We first sampled 100 proteins and calculated means square errors cover all pairs of the 100 proteins. The errors are baselines. For 3D-VQ-VAE, we calculated mean squared errors between original vMaSIF and reconstructed vMaSIF calculated by 3D-VQ-VAE over all proteins from CATH 4.2 test set.

| Model | SDF | Charge | Hydrophobicity | Hydrogen bond potential |
| --- | --- | --- | --- | --- |
| Dataset | 3.9e-4 | 2.6e-2 | 6.6 | 9.2e-1 |
| 3D-VQ-VAE | 4.9e-6 | 2.1e-4 | 1.0e-2 | 1.0e-2 |

To develop a protein surface generation model, we first need to develop a model that approximate the distribution of protein surfaces. Our proposed model, ShapeProt, achieved the goal by utilizing 3D-VQVAE. 3D-VQVAE capture the distribution of protein surfaces, and has a latent space capable of encompassing the protein surface distribution. Since our final goal is to perform generative surface modification in 3D space, spatial consistency between the latent and real world allows us to conduct the surgery more intuitively. Because the 3D VQVAE incorporates 3D CNN-based architectures and a 3D grid latent space, we earned the continuous preservation of spatial inductive bias throughout the whole networks, guaranteeing the spatial consistency for ShapeProt. After examining the visualization analysis, we confirmed that the latent space’s grid space precisely matches the corresponding locations in the real space. In addition, the utilization of VQVAE effectively avoids the issue of Posterior Collapse that is commonly encountered in conventional VAEs, hence ensuring stable training.

The encoder comprises a three-dimensional convolutional neural network (3D CNN) to represent the four-channeled voxelized MaSIF because grid-patch-based convolution modeling is suitable for accurately representing the localized environment of protein molecular surfaces. Furthermore, including residuals enables continuous information delivery throughout the whole network. Like the Unet encoder architecture, as the layers deepen, the grid size is reduced by half while the number of channels is doubled. Multi-channeld N-sized grid features are obtained through the encoder. In this work, we set N as 24.

Then, the grid features from the encoder are transformed into a three-channel latent grid via a single 3D CNN layer before latent feature computation. After that, these latent grid features are re-represented using a vector quantizer. The vector quantizer expresses each grid feature with the corresponding feature of the trainable codebook based on the distance-based similarity. To enhance the expressiveness of our VQ-VAE, we adopted two strategies. First, we employed distinct vector quantizers for each of the four MaSIF channels. The analysis showed apparent differences in the distributions of these four characteristics, with just a few noticeable relationships (see Figure 6).

Consequently, we recognized the difficulty of effectively capturing the four diverse attributes with a single codebook. Also, empirical findings demonstrated that employing separate vector quantizers for these four distinct features resulted in the best performance (See Figure 7. The second strategy is to use residual quantization. RQ-VAE [49] proved that more than multiple quantization allowed fluent representation, alleviating the inevitable information loss during compression using vector quantization. This method repeated codebook feature searches and accumulated them through simple summation, allowing for richer information than conventional VQ-VAE. The residual quantization not only maintains superior quality representation but also operates with remarkable computing efficiency. Subsequently, to generate input in the decoder, we concatenate the four quantized features channel-wise. These combined features are then passed to a 3D CNN layer, facilitating the amalgamation of the four distinct features and yielding the decoder input grid features.

In our decoder architecture, four separate decoders are employed. Although predicting all four features in a single decoder with a four-channel might be more efficient, our experiments demonstrated that this strategy did not show the best reconstruction performance (see Supplementary Material). We argue that this result is due to the difference between the four features. Consequently, we developed individual decoders for each feature, allowing each decoder to focus on its respective information. Notably, since we utilize a single encoder until the latent stage, the learned latent representation encompasses all four features concurrently. Additionally, decoding occurs after separate quantization on the latent features and subsequent integration of the four features; thus, decoding of each feature is correlated. To merge the reconstructed four features, we design a novel Molecular surface Sketch and Chemical Coloring (MSCC) process. We first define the molecular surfaces based on the decoded SDF, followed by marching cubes [50] (Molecular surface sketch). Then we calculate and assign the Gaussian-weighted average of chemical features from the top K nearest voxel grid points for the each vertices of the reconstructed mesh (Chemical Coloring). The whole architecture is depicted in Figure 2 (a).

**Figure 2:**
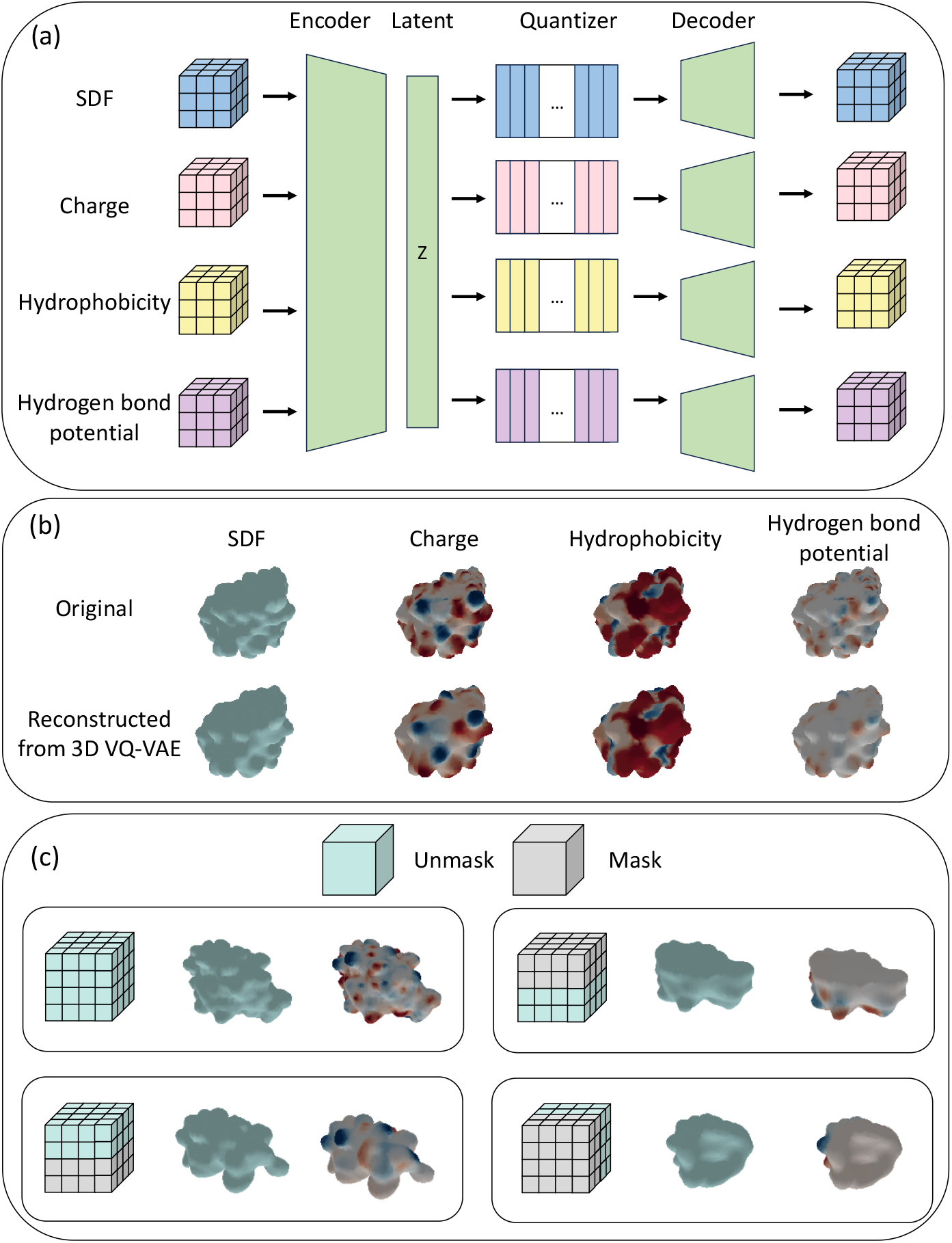
(a), an illustration of 3D-VQVAE. (b), Examples showing reconstruction quality of the trained 3D-VQVAE. (c), our 3D-VQVAE has a spatial consistency between latent grids and grids of vMaSIF in real space.

Figure 2 (b) illustrate the reconstruction ability of trained 3D-VQVAE. It is worth mentioning that 3D-VQVAE preserves complicated information although the protein surfaces exhibit significant irregularity and possess high-frequency properties.

Figure 2 (c) shows the spacial consistency between the latent grids and output grids in real space. First, We masked out specific grids of a latent grid of a protein sample along various views (top, bottom and front). Then, the masked latent grids are fed into the decoder and vMaSIFs are calculated. Each vMaSIF misses the some parts of the original vMaSIF according to the used mask.

### 2.3 Latent diffusion modeling on shape latent space

We train a latent diffusion model upon latent representation z obtained by the trained encoder. We employ a time-conditional 3D UNet [51] for this purpose. The optimization of the diffusion model is carried out by employing a simplified denoising objective [14]. During the process of diffusion inference, the latent codes are acquired by gradually removing noise that is sampled from a standard normal distribution. The code is subsequently inputted into distinct decoders, leading to the successful decoding of a voxel grid consisting of four channels. Additionally, we apply conditional generation via classifier-free guidance [40] to control the learned distribution by conditional inputs and cross-attention mechanisms. Because our goal is to obtain various sequences that matches diverse generated shapes while maintaining overall structure of a given protein, backbone condition is employed, which enable the shape diffusion model to generate surfaces with similar backbone. GVP [52] encoders are employed to extract conditional embeddings of backbone. Also, we utilized mask-based inpainting [53]. Figure 3 (a) depicts the diffusion networks and conditioning methods.

**Figure 3:**
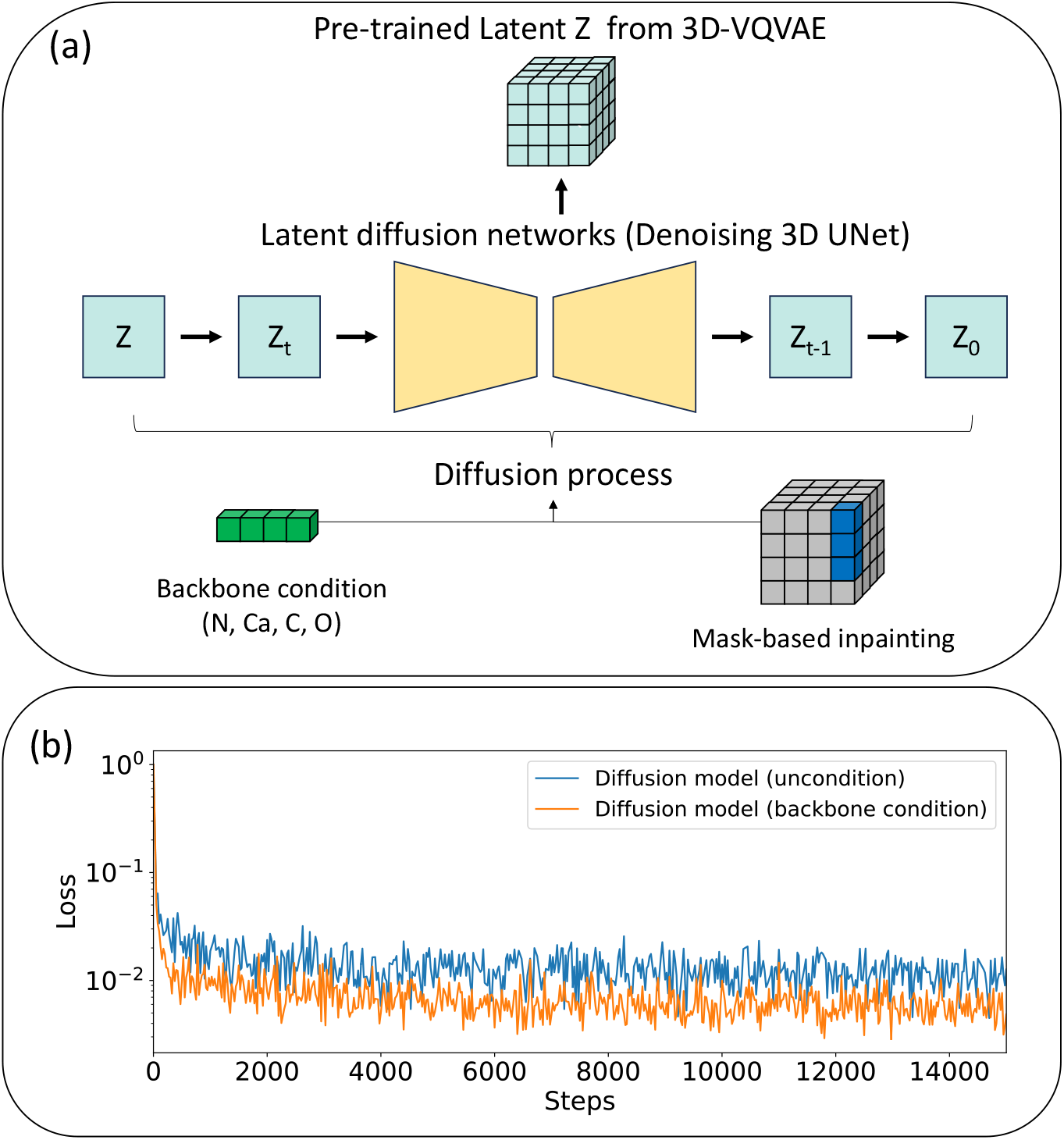
(a), an illustration describing our latent diffusion models. We adopted conditioning method based on classifer-free-guidance and mask-based inpainting. (b), training curves of diffusion model with and without backbone condition.

**Figure 4:**
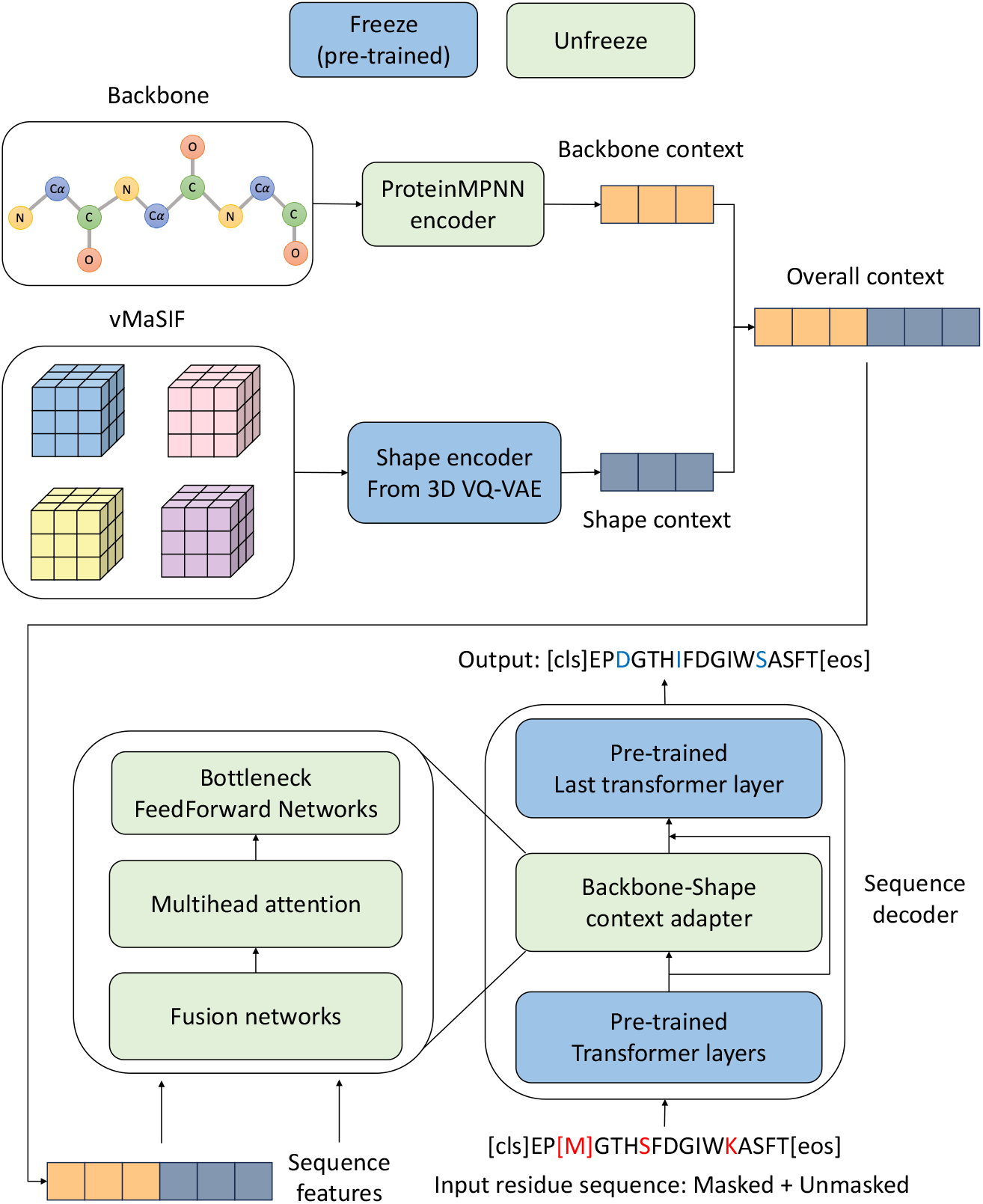
Overview of inverse-shape model scheme. Our proposed Surf-Design model conditions on both backbone and shape contexts.

**Figure 5:**
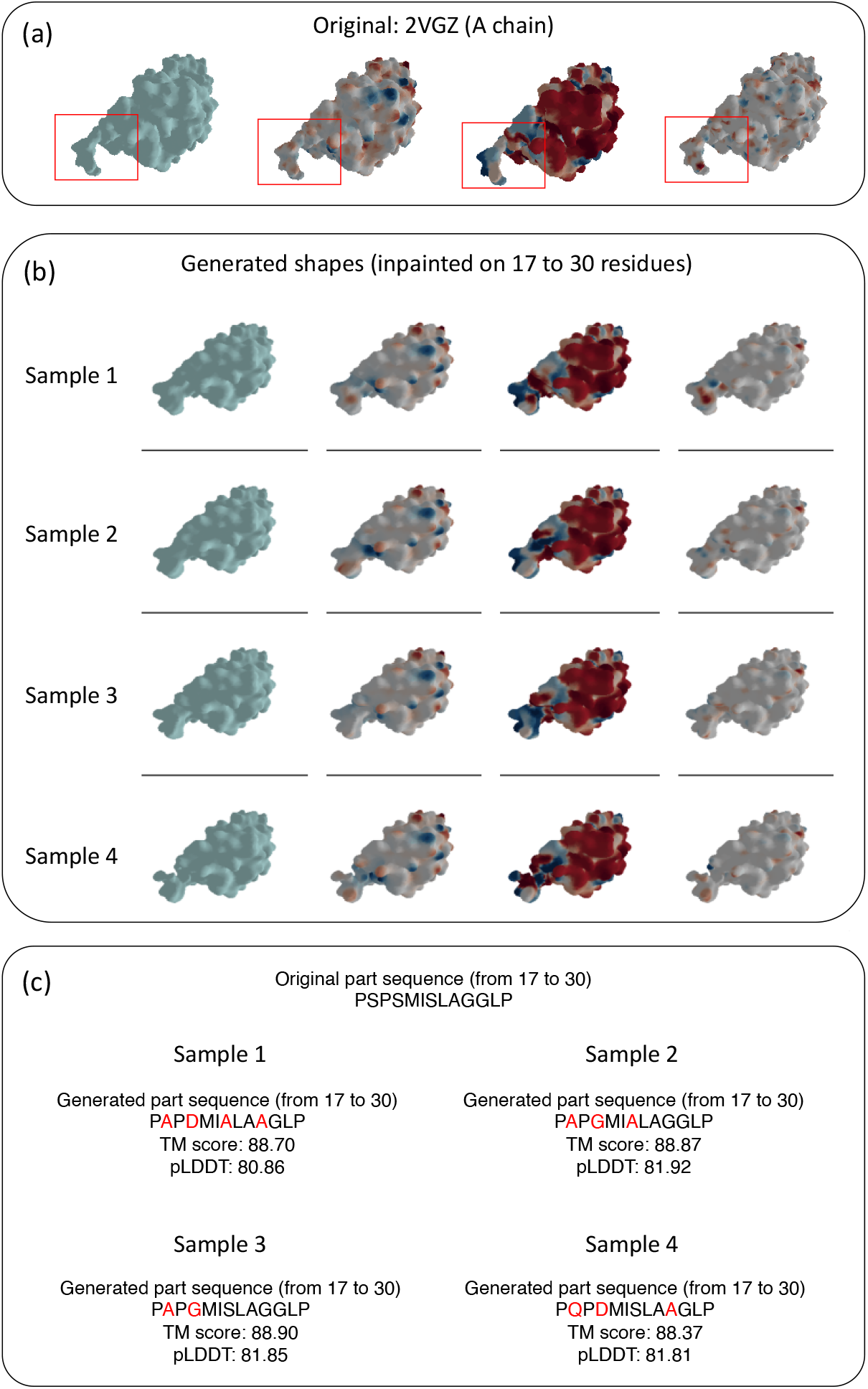
We conducted shape surgery using our ShapeProt framework. (a) We selected subregion of a protein (PDB ID: 2VGZ), (b) generated diverse shape samples with inpainting and conditioning, and (c) finally traslated the shapes into sequences and calculated various metrics.

### 2.4 Inverse shape approach

Inverse folding is a protein design strategy that focuses on the generation of promising sequences while preserving a given backbone structure [30, 52, 54, 55]. The fixed backbone design is grounded in structure consistency so that inverse folding enables investigating diverse sequence spaces while conserving the original protein’s function. Since protein functionality is closely related to the surface, leveraging additional context related to the surface would be beneficial for fine-grained protein design. This concept has some similarities to contemporary top-down methodologies that employ structural constraints for protein design. The surfaces generated from backbone conditioning on ProMOL-E represent surface candidates for a given fixed backbone. Therefore, we can translate those surfaces into the corresponding possible sequences that match the generated surface by adding surface context to inverse folding.

To validate this concept, we extend the LM-Design framework by adding surface context, and we call the model Surf-Design. LM-design is a state-of-the-art inverse-folding model that generates sequences using a pre-trained protein language model adapted by structural context. We conducted training using the CATH 4.2 dataset [56] and evaluated the two models based on sequence recovery for a fair comparison. In this setting, Surf-Design and LM-design show similar performances of 50.40 % and 51.11 %, respectively, on test set of CATH 4.2. Interestly, as shown in Figure 8 the generated sequences obtained from between LM-Design and Surf-Design are quite different, suggesting that shape contexts gave different context to sequence decoder. We call the approach inverse shape.

**Figure 6:**
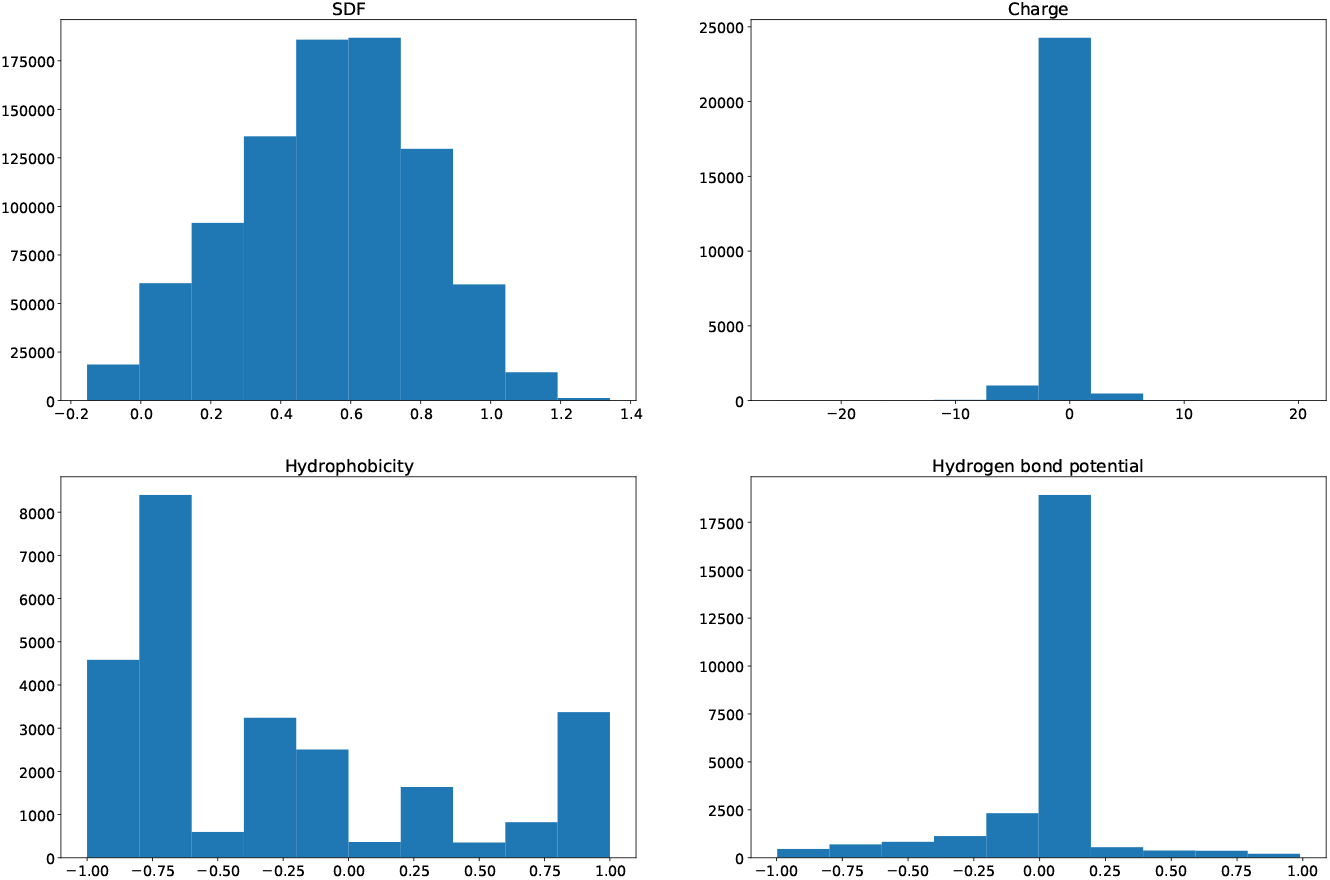
Distribution of SDF, charge, hydrophobicity, and hydrogen bond potential of 3DZZ_A protein sample.

**Figure 7:**
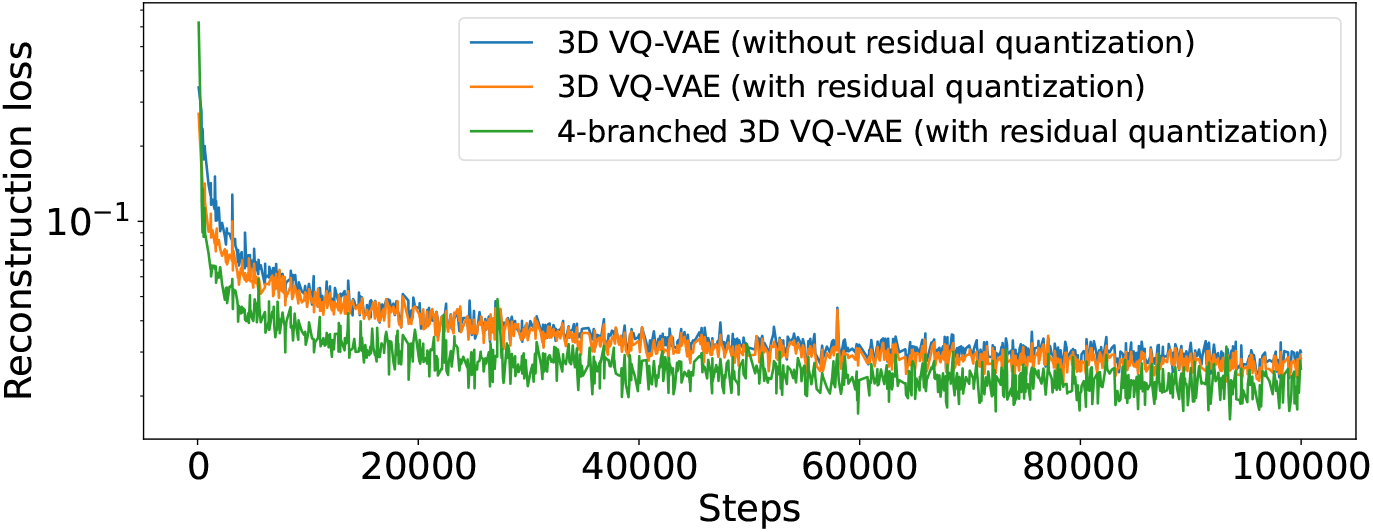
Reconstruction loss curves of variants of 3D-VQVAE.

**Figure 8:**
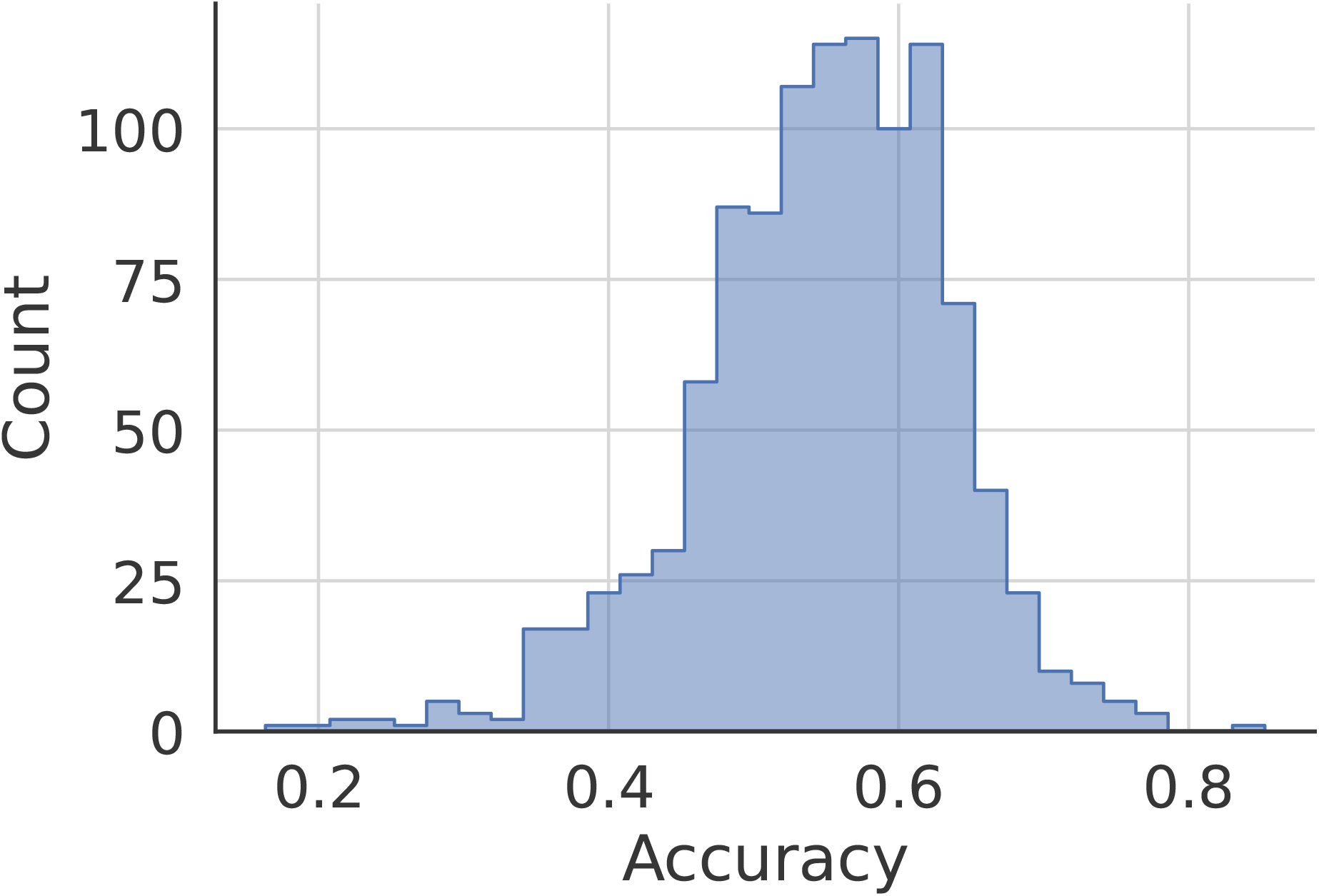
Difference between sequences from LM-Design and Surf-Design.

**Table 2:** Sequence recovery of various pLM-based sequence captioner on CATH4.2 test set.

| Model | Backbone | Surface | Sequence Recovery |
| --- | --- | --- | --- |
| LM-Design | ✓ |  | 51.11 |
| Surf-only-Design |  | ✓ | 7.10 |
| Surf-Design | ✓ | ✓ | 50.40 |

### 2.5 Diffusion-based Protein Shape Surgery and sequence design

Utilizing the three models above enables a distinctive top-down protein design approach, ShapeProt, where the diverse shapes for specific regions are generated, and the subsequent generation of corresponding sequences is conducted. To validate this, we conducted a case study on the A chain of the 2VGZ protein. We selected the surface region consisting of residues 17-30 from the A chain. Initially, we computed the latent z of the A chain using 3D-VQ-VAE and calculated the backbone condition of the A chain using the backbone encoder. Latent grid parts corresponding to this surface region were then determined based on Euclidean distances. Afterward, diverse surfaces were generated using DDIM sampling. The utilization of backbone conditioning results in the generated surfaces having similar backbones. Moreover, during diffusion sampling, latent grid parts corresponding to the surface region remained, and the other parts were replaced by the transition state corresponding to the original latent z, ensuring generation specifically for the target surface while preserving the rest as much as possible. Figure 5 illustrates the generation of different shapes for the target surfaces highlighted by the red box (see Figure 5a). Subsequently, using Surf-Design, inpainting sequence generation was performed for the target surfaces, yielding the final sequences (see Figure 5c). Interestingly, we can see that there are variations in amino acids compared to the original sequence part with regard to the generated shapes. Also, we conducted structure prediction using ESMFold for the generated sequences, achieving a high TM-score of 88.

## 3 Discussion

Protein design aims to generate and manipulate proteins with functionalities tailored for specific applications, and the function is strongly tied to the geometric and chemical features of functional sites on the protein surface. The conventional protein sequence design methods predominantly concentrate on designing proteins based on either sequence or geometric structure, but we assumed that this design is equivalent to the design of the protein surface. Given the abstract and complex 3D nature of protein surfaces, traditional approaches using 3D structure geometric learning models face challenges in modeling protein shapes. In this study, we successfully modeled such surfaces using SDF-based 3D deep learning, developing a 3D-VQ-VAE capable of capturing 3D geometric and chemical surfaces. We have developed a new type of protein generative modeling way to generate a variety of target surfaces by integrating advanced generative models, the latent diffusion model, with conditioning and inpainting techniques. Recognizing the indispensability of sequences even for novel surfaces, we formulated the surface-to-sequence captioning problem. We successfully addressed the task by developing a surface-conditioned inverse-folding model that could generate various sequences when given backbone and surface context. Our case study demonstrated the ability of our protein design framework, ShapeProt, to perform protein shape surgery by combining three models. Our work introduces a pioneering approach to surgery on 3D protein shapes directly and designing sequences based on these shapes. We hope our work showcases the applicability of 3D deep learning modeling in the emerging field of top-down protein design.

## 4 Methods

### 4.1 Data preparation

#### vMaSIF generation

Initially, we calculated MaSIF data and then constructed a voxelized MaSIF database. The MaSIF data were processed using the identical parameters specified in the MaSIF publication. The specific procedures are as follows: Protonation of all proteins in the datasets was performed using the Reduce software [57], then triangulation was carried out using the MSMS program [43]. The triangulation process used a density of 3.0 and a water probe radius of 1.5 Å. Afterward, the protein meshes underwent downsampling and regularization to have a resolution of 1.0 Å, using the Pymesh [58]. The generated molecular mesh was centered using the mean coordinates for each axis and normalized by dividing the XYZ coordinates by 64. Subsequently, a 3D grid of coordinates with a size of 96×96×96 in the (−1, 1) range was created. SDF grids were then generated based on the distance from the molecular mesh vertices. We calculated chemical features of the protein meshes as follows.

#### Chemical feature calculation

To calculate hydrogen bond potential features, we first identified whether each vertex on the molecular surface is a potential donor or acceptor in hydrogen bonds based on their closest atom among polar hydrogen, nitrogen, or oxygen using hydrogen potential [59]. Afterward, a value from a Gaussian distribution was assigned to each vertex, considering the orientation between heavy atoms. The values varied from -1 (ideal position for a hydrogen bond acceptor) to +1 (ideal position for a hydrogen bond donor). To compute electronic charge features, we first prepared protein files using PDB2PQR [60], and APBS (v.1.5) [61] was utilized to calculate Poisson Boltzmann electrostatics for each protein. The charge at each vertex of the meshed surface was determined using Multivalue from the APBS suite. We assigned a hydropathy scalar value to each vertex based on the Kyte and Doolittle [62] scale of the amino acid identity of the atom closest to the vertex. The original range of these values was from -4.5 (the most hydrophilic) to +4.5 (the most hydrophobic). These values were later normalized to a normalized range between -1 and 1. For training purposes, a range of -3 to 3 was used for all features, obtained by multiplying the range of -1 to 1 by 3.

### 4.2 Model architecture

#### 3D VQ-VAE

At first, a four-channel vMaSIF input undergoes 3D convolution, resulting in 64-channeled grid features before passing through encoder part. The encoder comprises three encoding blocks, two mid-blocks, and one attention block. Each encoding block and mid-block consists of two subblocks composed of a group normalization layer, a sigmoid-based nonlinear layer, and a 3D convolution layer. In the encoding block, the features are updated through two subblocks, and a residual connection is applied at the end. The grid resolution decreases by a factor of 2 due to convolution-based downsampling after each passage through the encoding block. The updated features from the encoding block are then updated in the first mid-block and processed through an attention-based block. The attention block employs convolution-based query, key, and value attention. Subsequently, the features are updated in the second mid-block, followed by normalization and nonlinear operation. Finally, the features are projected to a latent channel of 3 before quantization. The convolution layers used in the subblock and attention block have a kernel size of 3, stride of 1, and padding of 1.

The quantizer consists of an 8192-sized codebook and three features represent each code. We employ residual quantization, which involves four rounds to provide codebook-based grid features. Each grid feature is represented by the closest code feature based on inter-feature distances in each iteration. Subsequently, we subtracted the codebook-based grid feature from the original grid feature to avoid duplicated information accumulation. After conducting the same process four times, the four codebook-based grid features obtained are summed to yield the final quantized grid feature. Meanwhile, a quantizer is created for each SDF, hydrophobicity, hydrogen bond potential, and charge. The four quantized grid features, which are encoded as codebook features, are updated through a block comprising a 3D convolution layer, normalization layer, GELU layer [63], and 3D convolution layer in order.

The decoder comprises three decoding blocks, two mid-blocks, and one attention block, and each block has the same architecture as the encoding block, mid-block, and attention block used in the encoder, respectively. Initially, the quantized feature undergoes a single convolution layer to expand the channels from 3 to 64. Subsequently, it gets updated in sequence through the first mid-block, attention block, and second mid-block. Subsequently, the feature pass through the decoding blocks, and upsampling operations are applied, resulting in a doubling of resolution at each block. Afterward, the features pass update networks consisting of two linear layers and two GELU layers. Finally, the output goes through the final linear layer, resulting in a single-channeled output. There are four decoders corresponding to the four quantizers. The grid outputs produced from each decoder are combined channel-wise to construct the final reconstructed vMaSIF with four channels.

#### Diffusion model

The conventional latent diffusion model processes 2D latent feature maps using 2D convolutions [16]. In our approach, we utilized a variant employing 3D convolutions to handle a 3D grid. The overall structure follows a Unet architecture [64], consisting of three parts: input block, middle block, and output block. The Input block comprises three subblocks, each consisting of two residual blocks. Spatial transformer blocks are appended to the second and third blocks. The channel size increases by a factor of 1, 2, and 3 after passing through each subblock. The middle block consists of a residual block, a spatial transformer block, and another residual block in sequence. The output block is the same as the input block but adopts upsampling operations. The channel size undergoes a multiplicative decrease of 1, 2, and 3 after traversing each subblock. The residual block layers take the latent grid feature and time step embedding as input, integrating them using multiple convolution layers to update and integrate the latent grid feature and time step information. The spatial transformer block utilizes cross attention with conditioning context as input, injecting context into the latent grid feature. Latent diffusion is applied to 3-channel grid features just before the 3D VQ-VAE quantization, so the 3-channel grid features are expanded to 224 channels using a 3D convolution layer and then processed through the aforementioned Unet-like main networks. Finally, a 3D convolution layer is applied to the output from Unet, yielding a 3-channel grid output used in denoising training.

#### Surf-Design

The Surf-Design framework is built upon the ProteinMPNN-LM-Design model [65] and consists of four main components: structure encoder, shape encoder, sequence decoder, and shape-backbone context adapter. The structure encoder employs the published version of ProteinMPNN [19], and the sequence decoder utilizes the pre-trained ESM-1b 650M. The Shape Encoder consists of subnetworks of the pre-trained 3D-VQ-VAE encoder, from the opening to the convolution layer, which is just before the quantization part in the pre-trained 3D-VQ-VAE. The pre-trained sequence decoder and shape encoder are frozen without updates, while the remaining structure encoder and adapter parameters are updated. Initially, the structure encoder extracts backbone context from the backbone, and the shape encoder calculates shape context from vMaSIF. The two contexts are combined to form an overall context, which is then updated through self-attention using a standard transformer layer to integrate the two information. Subsequently, the sequence features are updated via the transformer layers of the sequence decoder until the last transformer layer. Then, the adapter operation is applied, where sequence features act as queries and undergoes cross-attention with the overall context serving as keys and values. Finally, the transformed sequence features pass through the last transformer layer for token prediction.

### 4.3 Model training

#### 3D VQ-VAE

The 3D VQ-VAE is trained to quantize and reconstruct vMaSIF. A vMaSIF **X** is encoded by the encoder **ENC** to obtain a feature grid 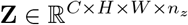, which is then represented as a quantized stacked grid 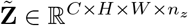 based on a stacked grids of codes **M** ϵ [**K**]^*C×H×W ×D*^ via residual quantization with depth *D* as follows:

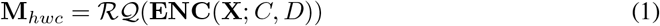

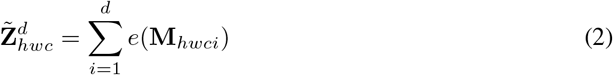

where *e, ℛ𝒬*, and 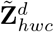 denote a code embedding layer, residual quantization operation, and the quantized feature grid at depth *d*, respectively. Subsequently, 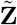 undergoes reconstruction through the decoder **DEC** as 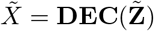. The training of the 3D VQVAE involves two types of losses: the reconstruction loss *L*_*recon*_ and the Commitment loss *ℒ*_*commit*_ as follows:

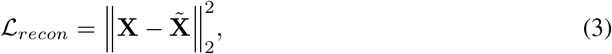

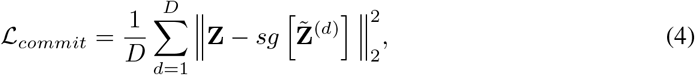

where *sg* [·] denote the stop-gradient operator. The straight-through estimator [39] is utilized for the backpropagation through the residual quantizer layer. Since there are four quantizer in 3D-VQVAE, total training loss ℒ_*total*_ for 3D-VQVAE is as follows:

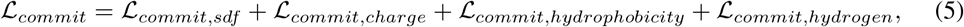

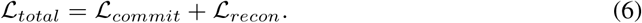

#### Diffusion model

Diffusion models (DMs) [66, 14] are probabilistic models that learn to acquire a data distribution by denoising a Gaussian variable. DMs define two distinct processes, (1) forward and (2) reverse processes, operating in reverse directions. In the forward process *q*(*x*_0:*T*_), a data sample **x** is gradually transformed to pure Gaussian noise. This diffusion process is defined as a Markov chain as follows:

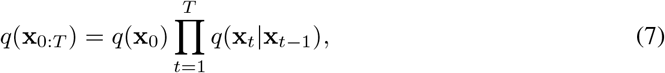

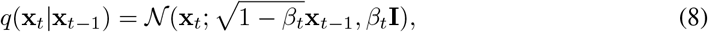

where *β*_*t*_ is the noise scheduling hyperparameter. Following the reparameterization trick [14], we can sample **x** at an arbitrary time step *t* in the forward process:

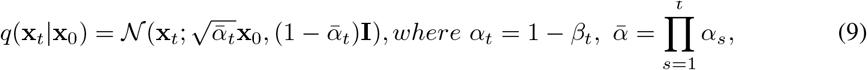

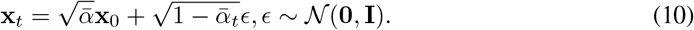

In reverse process *p*_*θ*_(*x*_0:*T*_, a data sample is recovered by gradually removing noise from pure Gaussian. This reverse process is defined as a Markov chain and learned with parameter *θ*:

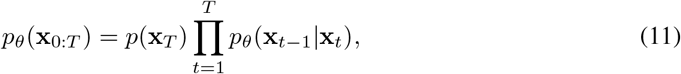

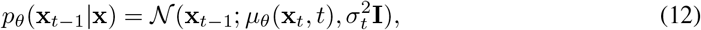

where, *p*(**x**_*T*_) = 𝒩(**x**_*T*_ ; **0, I**) indicates the standard Gaussian distribution, *µ*_*θ*_(**x**_*t*_, *t*) denotes a denoising network parameterized by *θ*,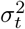 is a time-step dependant variance. We can sample **x**_*t−*1_ as:

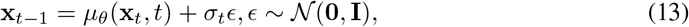

Following [14], *µ*_*θ*_(**x**_*t*_, *t*) is simplified:

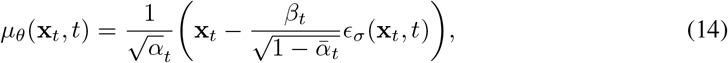

where *ϵ*_*σ*_(**x**_*t*_, *t*) is a neural network with parameter *θ*. We predict noises from noisy input *x*_*t*_ with the *ϵ*_*σ*_(**x**_*t*_, *t*) so that our training objective is mean squared error between the predicted noise *ϵ*_*θ*_(**x**_*t*_, *t*) and the added noise *ϵ* at a time step as follows:

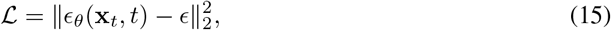

Meanwhile, we added backbone condition to the diffusion model, so, backbone condition *c*_*b*_ is fed into the diffusion process. Finally, the reverse process becomes *p*_*θ*_(**x**_*t −*1_ |**x**, *c*) and the final objective is as follows:

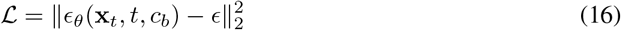

#### Surf-Design

Surf-Design utilized an MLM-based protein language model as a sequence decoder, following the approach proposed in LM-design. The pLMs is used in a non-autoregressive decoding fashion. This method is known as conditional masked language modeling (CMLM), which enables parallel decoding, enhancing the efficiency of conditional generation [67]. Additionally, the use of backbone and shape context condition-based adaptation allows the sequence decoder to perform backbone and shape-aware sequence captioning. Given a protein sample *P* with backbone *S* and vMaSIF *X*, the structure encoder computes the backbone context 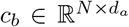, while the shape encoder computes shape context 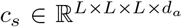. *N, L*, and *d*_*a*_ denote the length of sequence of protein *P*, the latent resolution of 3D-VQ-VAE encoder, and the hidden dimension of adapter. Then, the whole shape context 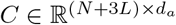 is obtained by concatenating the two context. Given sequence *R* = *R*_*masked*_ ∪ *R*_*observed*_ where *R*_*masked*_ and *R*_*observed*_ indicate the masked and visible residues respectively, the non-autoregressive process of Surf-Design is as follows:

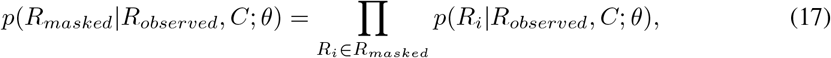

The *θ* is optimzed by minimizing the negative log-likelihood over every residue token in *R*_*masked*_.

**A Appendix**

## References

[1] Longxing Cao, Inna Goreshnik, Brian Coventry, James Brett Case, Lauren M. Miller, Lisa Kozodoy, Rita E. Chen, Lauren P. Carter, Alexandra C. Walls, Young-Jun Park, Eva-Maria Strauch, Lance J. Stewart, Michael Steven Diamond, David Veesler, and David Baker. De novo design of picomolar sars-cov-2 miniprotein inhibitors. Science (New York, N.y.), 370:426 – 431, 2020.

[2] Fabian Sesterhenn, Che Yang, Jaume Bonet, Johannes T Cramer, Xiaolin Wen, Yimeng Wang, Chi-I Chiang, Luciano A Abriata, Iga Kucharska, Giacomo Castoro, et al. De novo protein design enables the precise induction of rsv-neutralizing antibodies. Science, 368(6492):eaay5051, 2020.

[3] Daniel-Adriano Silva, Shawn Yu, Umut Y. Ulge, Jamie B. Spangler, Kevin M. Jude, Carlos Labão-Almeida, Lestat R. Ali, Alfredo Quijano-Rubio, Mikel Ruterbusch, Isabel Leung, Tamara Biary, Stephanie J. Crowley, Enrique Sánchez Marcos, Carl Walkey, Brian D. Weitzner, Fátima Pardo-Avila, Javier Castellanos, Lauren P. Carter, Lance J. Stewart, Stanley R. Riddell, Marion Pepper, Gonçalo J. L. Bernardes, Michael L Dougan, K. Christopher Garcia, and David Baker. De novo design of potent and selective mimics of il-2 and il-15. Nature, 565:186 – 191, 2019.

[4] Jessica Marcandalli, Brooke Fiala, Sebastian Ols, Michela Perotti, Willem de van der Schueren, Joost Snijder, Edgar A Hodge, Mark A. Benhaim, Rashmi Ravichandran, Lauren P. Carter, William Sheffler, Livia Brunner, Maria Lawrenz, Patrice M. Dubois, Antonio Lanzavecchia, Federica Sallusto, Kelly K. Lee, David Veesler, Colin E. Correnti, Lance J. Stewart, David Baker, Karin Loré, Laurent Perez, and Neil P. King. Induction of potent neutralizing antibody responses by a designed protein nanoparticle vaccine for respiratory syncytial virus. Cell, 176:1420 – 1431.e17, 2019.

[5] Sarel Jacob Fleishman, Timothy A. Whitehead, Damian C. Ekiert, Cyrille Dreyfus, Jacob E. Corn, Eva-Maria Strauch, Ian A. Wilson, and David Baker. Computational design of proteins targeting the conserved stem region of influenza hemagglutinin. Science, 332:816 – 821, 2011.

[6] Greta Giordano-Attianese, Pablo Gainza, Elise F Gray-Gaillard, Elisabetta Cribioli, Sailan Shui, Seong Hoon Kim, Mi-Jeong Kwak, Sabrina S. Vollers, Angel De Jesus Corria Osorio, Patrick Reichenbach, Jaume Bonet, Byung-Ha Oh, Melita Irving, George Coukos, and Bruno E. Correia. A computationally designed chimeric antigen receptor provides a small-molecule safety switch for t-cell therapy. Nature Biotechnology, 38:426 – 432, 2020.

[7] Longxing Cao, Brian Coventry, Inna Goreshnik, Buwei Huang, William Sheffler, Joon Sung Park, Kevin M. Jude, Iva Markovći, Rameshwar U. Kadam, Koen H. G. Verschueren, Kenneth Verstraete, Scott Thomas Russell Walsh, Nathaniel R. Bennett, Ashish Phal, Aerin Yang, Lisa Kozodoy, Michelle DeWitt, Lora K. Picton, Lauren M. Miller, Eva-Maria Strauch, Nic Debouver, Allison Pires, Asim K. Bera, Samer Halabiya, Bradley Hammerson, Wei Yang, Steffen M. Bernard, Lance J. Stewart, Ian A. Wilson, Hannele Ruohola-Baker, Joseph Schlessinger, Sangwon Lee, Savvas N. Savvides, K. Christopher Garcia, and David Baker. Design of protein-binding proteins from the target structure alone. Nature, 605:551 – 560, 2022.

[8] Isaac D. Lutz, Shunzhi Wang, Christoffer H Norn, Alexis Courbet, Andrew J. Borst, Yan Ting Zhao, Annie M. Dosey, Longxing Cao, Jinwei Xu, Elizabeth M. Leaf, Catherine Treichel, Patrisia Litvicov, Zhe Li, Alexander D. Goodson, Paula Rivera-Sánchez, Ana - Maria Bratovianu, Minkyung Baek, Neil P. King, Hannele Ruohola-Baker, and David Baker. Top-down design of protein architectures with reinforcement learning. Science, 380:266 – 273, 2023.

[9] John B Ingraham, Max Baranov, Zak Costello, Karl W Barber, Wujie Wang, Ahmed Ismail, Vincent Frappier, Dana M Lord, Christopher Ng-Thow-Hing, Erik R Van Vlack, et al. Illuminating protein space with a programmable generative model. Nature, pages 1–9, 2023.

[10] Cyrus Chothia and Joël Janin. Principles of protein–protein recognition. Nature, 256:705–708, 1975.

[11] Loredana Lo Conte, Cyrus Chothia, and Joël Janin. The atomic structure of protein-protein recognition sites. Journal of molecular biology, 285 5:2177–98, 1999.

[12] Tom B. Brown, Benjamin Mann, Nick Ryder, Melanie Subbiah, Jared Kaplan, Prafulla Dhariwal, Arvind Neelakantan, Pranav Shyam, Girish Sastry, Amanda Askell, Sandhini Agarwal, Ariel Herbert-Voss, Gretchen Krueger, T. J. Henighan, Rewon Child, Aditya Ramesh, Daniel M. Ziegler, Jeff Wu, Clemens Winter, Christopher Hesse, Mark Chen, Eric Sigler, Mateusz Litwin, Scott Gray, Benjamin Chess, Jack Clark, Christopher Berner, Sam McCandlish, Alec Radford, Ilya Sutskever, and Dario Amodei. Language models are few-shot learners. ArXiv, abs/2005.14165, 2020.

[13] Long Ouyang, Jeff Wu, Xu Jiang, Diogo Almeida, Carroll L. Wainwright, Pamela Mishkin, Chong Zhang, Sandhini Agarwal, Katarina Slama, Alex Ray, John Schulman, Jacob Hilton, Fraser Kelton, Luke E. Miller, Maddie Simens, Amanda Askell, Peter Welinder, Paul Francis Christiano, Jan Leike, and Ryan J. Lowe. Training language models to follow instructions with human feedback. ArXiv, abs/2203.02155, 2022.

[14] Jonathan Ho, Ajay Jain, and Pieter Abbeel. Denoising diffusion probabilistic models. Advances in neural information processing systems, 33:6840–6851, 2020.

[15] Aditya Ramesh, Prafulla Dhariwal, Alex Nichol, Casey Chu, and Mark Chen. Hierarchical text-conditional image generation with clip latents. ArXiv, abs/2204.06125, 2022.

[16] Robin Rombach, Andreas Blattmann, Dominik Lorenz, Patrick Esser, and Björn Ommer. High-resolution image synthesis with latent diffusion models. In Proceedings of the IEEE/CVF conference on computer vision and pattern recognition, pages 10684–10695, 2022.

[17] Ting-Chun Wang, Ming-Yu Liu, Jun-Yan Zhu, Guilin Liu, Andrew Tao, Jan Kautz, and Bryan Catanzaro. Video-to-video synthesis. In Neural Information Processing Systems, 2018.

[18] Patrick Esser, Johnathan Chiu, Parmida Atighehchian, Jonathan Granskog, and Anastasis Germanidis. Structure and content-guided video synthesis with diffusion models. ArXiv, abs/2302.03011, 2023.

[19] Justas Dauparas, Ivan V. Anishchenko, Nathaniel R. Bennett, Hua Bai, Robert J. Ragotte, Lukas F. Milles, Basile I. M. Wicky, Alexis Courbet, Robbert J. de Haas, Neville P. Bethel, Philip J. Y. Leung, Timothy F. Huddy, Samuel J. Pellock, Doug K Tischer, F. Chan, Brian Koepnick, Hao A Nguyen, Alex Kang, Banumathi Sankaran, A. K. Bera, Neil P. King, and David Baker. Robust deep learning based protein sequence design using proteinmpnn. Science (New York, N.Y.), 378:49 – 56, 2022.

[20] Jacob Devlin, Ming-Wei Chang, Kenton Lee, and Kristina Toutanova. Bert: Pre-training of deep bidirectional transformers for language understanding. In North American Chapter of the Association for Computational Linguistics, 2019.

[21] Alec Radford and Karthik Narasimhan. Improving language understanding by generative pre-training. 2018.

[22] Mohammad Bavarian, Heewoo Jun, Nikolas Tezak, John Schulman, Christine McLeavey, Jerry Tworek, and Mark Chen. Efficient training of language models to fill in the middle. arXiv preprint arXiv:2207.14255, 2022.

[23] Alexander Rives, Siddharth Goyal, Joshua Meier, Demi Guo, Myle Ott, C. Lawrence Zitnick, Jerry Ma, and Rob Fergus. Biological structure and function emerge from scaling unsupervised learning to 250 million protein sequences. Proceedings of the National Academy of Sciences of the United States of America, 118, 2019.

[24] Noelia Ferruz, Steffen Schmidt, and Birte Höcker. Protgpt2 is a deep unsupervised language model for protein design. Nature Communications, 13, 2022.

[25] Ali Madani, Ben Krause, Eric R. Greene, Subu Subramanian, Benjamin P. Mohr, James M. Holton, Jose Luis Olmos, Caiming Xiong, Zachary Z Sun, Richard Socher, James S. Fraser, and Nikhil Vijay Naik. Large language models generate functional protein sequences across diverse families. Nature Biotechnology, pages 1–8, 2023.

[26] Youhan Lee and Hasun Yu. Protfim: Fill-in-middle protein sequence design via protein language models. arXiv preprint arXiv:2303.16452, 2023.

[27] Marjan Ghazvininejad, Omer Levy, Yinhan Liu, and Luke Zettlemoyer. Mask-predict: Parallel decoding of conditional masked language models. In Conference on Empirical Methods in Natural Language Processing, 2019.

[28] Jin Su, Chenchen Han, Yuyang Zhou, Junjie Shan, Xibin Zhou, and Fajie Yuan. Saprot: Protein language modeling with structure-aware vocabulary. bioRxiv, pages 2023–10, 2023.

[29] Michel van Kempen, Stephanie S Kim, Charlotte Tumescheit, Milot Mirdita, Cameron LM Gilchrist, Johannes Söding, and Martin Steinegger. Foldseek: fast and accurate protein structure search. Biorxiv, pages 2022–02, 2022.

[30] John Ingraham, Vikas K. Garg, Regina Barzilay, and T. Jaakkola. Generative models for graph-based protein design. In DGS@ICLR, 2019.

[31] Namrata Anand, Raphael R. Eguchi, Irimpan I. Mathews, Carla P. Perez, Alexander Derry, Russ B. Altman, and Po-Ssu Huang. Protein sequence design with a learned potential. Nature Communications, 13, 2022.

[32] Chloe Hsu, Robert Verkuil, Jason Liu, Zeming Lin, Brian L. Hie, Tom Sercu, Adam Lerer, and Alexander Rives. Learning inverse folding from millions of predicted structures. bioRxiv, 2022.

[33] Wengong Jin, Jeremy Wohlwend, Regina Barzilay, and T. Jaakkola. Iterative refinement graph neural network for antibody sequence-structure co-design. ArXiv, abs/2110.04624, 2021.

[34] Jeong Joon Park, Peter Florence, Julian Straub, Richard Newcombe, and Steven Lovegrove. Deepsdf: Learning continuous signed distance functions for shape representation. In Proceedings of the IEEE/CVF conference on computer vision and pattern recognition, pages 165–174, 2019.

[35] Ben Mildenhall, Pratul P Srinivasan, Matthew Tancik, Jonathan T Barron, Ravi Ramamoorthi, and Ren Ng. Nerf: Representing scenes as neural radiance fields for view synthesis. Communications of the ACM, 65(1):99–106, 2021.

[36] Heewoo Jun and Alex Nichol. Shap-e: Generating conditional 3d implicit functions. arXiv preprint arXiv:2305.02463, 2023.

[37] Jaehyeok Shim, Changwoo Kang, and Kyungdon Joo. Diffusion-based signed distance fields for 3d shape generation. In Proceedings of the IEEE/CVF Conference on Computer Vision and Pattern Recognition, pages 20887–20897, 2023.

[38] Yen-Chi Cheng, Hsin-Ying Lee, Sergey Tulyakov, Alexander G Schwing, and Liang-Yan Gui. Sdfusion: Multimodal 3d shape completion, reconstruction, and generation. In Proceedings of the IEEE/CVF Conference on Computer Vision and Pattern Recognition, pages 4456–4465, 2023.

[39] Aäron van den Oord, Oriol Vinyals, and Koray Kavukcuoglu. Neural discrete representation learning. ArXiv, abs/1711.00937, 2017.

[40] Jonathan Ho. Classifier-free diffusion guidance. ArXiv, abs/2207.12598, 2022.

[41] Pablo Gainza, Freyr Sverrisson, Frederico Monti, Emanuele Rodola, D Boscaini, MM Bronstein, and BE Correia. Deciphering interaction fingerprints from protein molecular surfaces using geometric deep learning. Nature Methods, 17(2):184–192, 2020.

[42] Michael L Connolly. Solvent-accessible surfaces of proteins and nucleic acids. Science, 221(4612):709–713, 1983.

[43] Michel F. Sanner, Arthur J. Olson, and Jean-Claude Spehner. Reduced surface: an efficient way to compute molecular surfaces. Biopolymers, 38 3:305–20, 1996.

[44] Jeong Joon Park, Peter R. Florence, Julian Straub, Richard A. Newcombe, and S. Lovegrove. Deepsdf: Learning continuous signed distance functions for shape representation. pages 165–174, 2019.

[45] Heewoo Jun and Alex Nichol. Shap-e: Generating conditional 3d implicit functions. ArXiv, abs/2305.02463, 2023.

[46] William E. Lorensen and Harvey E. Cline. Marching cubes: A high resolution 3d surface construction algorithm. Proceedings of the 14th annual conference on Computer graphics and interactive techniques, 1987.

[47] Chaoyu Quan and Benjamin Stamm. Mathematical analysis and calculation of molecular surfaces. J. Comput. Phys., 322:760–782, 2016.

[48] Alex Krizhevsky, Ilya Sutskever, and Geoffrey E. Hinton. Imagenet classification with deep convolutional neural networks. Communications of the ACM, 60:84 – 90, 2012.

[49] Doyup Lee, Chiheon Kim, Saehoon Kim, Minsu Cho, and Wook-Shin Han. Autoregressive image generation using residual quantization. pages 11513–11522, 2022.

[50] Evgeni Chernyaev. Marching cubes 33: Construction of topologically correct isosurfaces. Technical report, 1995.

[51] Olaf Ronneberger, Philipp Fischer, and Thomas Brox. U-net: Convolutional networks for biomedical image segmentation. ArXiv, abs/1505.04597, 2015.

[52] Bowen Jing, Stephan Eismann, Patricia Suriana, Raphael J. L. Townshend, and Ron O. Dror. Learning from protein structure with geometric vector perceptrons. ArXiv, abs/2009.01411, 2020.

[53] Andreas Lugmayr, Martin Danelljan, Andres Romero, Fisher Yu, Radu Timofte, and Luc Van Gool. Repaint: Inpainting using denoising diffusion probabilistic models. In Proceedings of the IEEE/CVF Conference on Computer Vision and Pattern Recognition, pages 11461–11471, 2022.

[54] Namrata Anand-Achim, Raphael R. Eguchi, Alexander Derry, Russ B. Altman, and Po-Ssu Huang. Protein sequence design with a learned potential. Nature Communications, 13, 2020.

[55] Zhangyang Gao, Cheng Tan, and Stan Z. Li. Pifold: Toward effective and efficient protein inverse folding. ArXiv, abs/2209.12643, 2022.

[56] Christine A. Orengo, A. D. Michie, Susan Jones, David C. Jones, Mark B. Swindells, and Janet M. Thornton. Cath–a hierarchic classification of protein domain structures. Structure, 5 8:1093–108, 1997.

[57] J Michael Word, Simon C Lovell, Jane S Richardson, and David C Richardson. Asparagine and glutamine: using hydrogen atom contacts in the choice of side-chain amide orientation. Journal of molecular biology, 285(4):1735–1747, 1999.

[58] Q Zhou. Pymesh—geometry processing library for python. 2020.

[59] Tanja Kortemme, Alexandre V Morozov, and David Baker. An orientation-dependent hydrogen bonding potential improves prediction of specificity and structure for proteins and protein– protein complexes. Journal of molecular biology, 326(4):1239–1259, 2003.

[60] Todd J Dolinsky, Paul Czodrowski, Hui Li, Jens E Nielsen, Jan H Jensen, Gerhard Klebe, and Nathan A Baker. Pdb2pqr: expanding and upgrading automated preparation of biomolecular structures for molecular simulations. Nucleic acids research, 35(suppl_2):W522–W525, 2007.

[61] Nathan A Baker, David Sept, Simpson Joseph, Michael J Holst, and J Andrew McCammon. Electrostatics of nanosystems: application to microtubules and the ribosome. Proceedings of the National Academy of Sciences, 98(18):10037–10041, 2001.

[62] Jack Kyte and Russell F Doolittle. A simple method for displaying the hydropathic character of a protein. Journal of molecular biology, 157(1):105–132, 1982.

[63] Dan Hendrycks and Kevin Gimpel. Gaussian error linear units (gelus). arXiv preprint arXiv:1606.08415, 2016.

[64] Olaf Ronneberger, Philipp Fischer, and Thomas Brox. U-net: Convolutional networks for biomedical image segmentation. In Medical Image Computing and Computer-Assisted Intervention–MICCAI 2015: 18th International Conference, Munich, Germany, October 5-9, 2015, Proceedings, Part III 18, pages 234–241. Springer, 2015.

[65] Zaixiang Zheng, Yifan Deng, Dongyu Xue, Yi Zhou, Fei Ye, and Quanquan Gu. Structure-informed language models are protein designers. bioRxiv, pages 2023–02, 2023.

[66] Jascha Sohl-Dickstein, Eric Weiss, Niru Maheswaranathan, and Surya Ganguli. Deep unsupervised learning using nonequilibrium thermodynamics. In International conference on machine learning, pages 2256–2265. PMLR, 2015.

[67] Marjan Ghazvininejad, Omer Levy, Yinhan Liu, and Luke Zettlemoyer. Mask-predict: Parallel decoding of conditional masked language models. arXiv preprint arXiv:1904.09324, 2019.

